# Serine-129 phosphorylation of α-synuclein is a trigger for physiologic protein-protein interactions and synaptic function

**DOI:** 10.1101/2022.12.22.521485

**Authors:** Leonardo A. Parra-Rivas, Kayalvizhi Madhivanan, Lina Wang, Nicholas P. Boyer, Dube Dheeraj Prakashchand, Brent D. Aulston, Donald P. Pizzo, Kristen Branes-Guerrero, Yong Tang, Utpal Das, David A. Scott, Padmini Rangamani, Subhojit Roy

## Abstract

Phosphorylation of α-synuclein at the Serine-129 site (α-syn Ser129P) is an established pathologic hallmark of synucleinopathies, and also a therapeutic target. In physiologic states, only a small fraction of total α-syn is phosphorylated at this site, and consequently, almost all studies to date have focused on putative pathologic roles of this post-translational modification. We noticed that unlike native (total) α-syn that is widely expressed throughout the brain, the overall pattern of α-syn Ser129P is restricted, suggesting intrinsic regulation and putative physiologic roles. Surprisingly, preventing phosphorylation at the Ser-129 site blocked the ability of α-syn to attenuate activity-dependent synaptic vesicle (SV) recycling – widely thought to reflect its normal function. Exploring mechanisms, we found that neuronal activity augments α-syn Ser-129P, and this phosphorylation is required for α-syn binding to VAMP2 and synapsin – two functional binding-partners that are necessary for α-syn function. AlphaFold2-driven modeling suggests a scenario where Ser129P induces conformational changes in the C-terminus that stabilizes this region and facilitates protein-protein interactions. Our experiments indicate that the pathology-associated Ser129P is an unexpected physiologic trigger of α-syn function, which has broad implications for pathophysiology and drug-development.

---

Aggregation of α-syn into Lewy bodies/neurites is a neuropathologic signature of Parkinson’s disease (PD) and related disorders; collectively called ‘synucleinopathies’. Almost all pathologically aggregated α-syn is phosphorylated at the Ser-129 position (Anderson et al., 2006), and many previous studies have explored the role of Ser129P in disease [reviewed in (Oueslati, 2016)]. Antibodies to Ser129 α-syn are the most sensitive marker for post-mortem neuropathologic diagnosis of synucleinopathies (Beach et al., 2008), and experimental studies routinely use this modification as a surrogate marker for pathology in animal models. Although only a small fraction (∼4%) of α-syn is phosphorylated at this site under normal conditions (Anderson et al., 2006), low-frequency events such as serine phosphorylation are known to regulate numerous biological properties (Nestler and Greengard, 1999). Indeed, one of the first studies mapping Ser129P in human α-syn (h-α-syn) noted its constitutive nature and suggested a physiologic role involving repeated phosphorylation and dephosphorylation (Okochi et al., 2000), nevertheless, almost all studies to date have examined α-syn Ser129 in the context of pathology. In most organisms, α-syn robustly targets to presynapses, though studies have also reported α-syn in the nucleus (Maroteaux et al., 1988; Pinho et al., 2019).

Examining localization of Ser129P α-syn, we found that the phosphorylated protein was detectable in presynapses and nuclei in mouse cultured hippocampal neurons (**Fig. 1A** and **Supp. Fig. 1A**). In mouse brains, unlike the broad distribution of native (total) α-syn, Ser129P α-syn was restricted to a subset of brain regions, with prominent synaptic and nuclear staining in the superficial cortex, hippocampus, olfactory and dopaminergic regions (**Fig. 1B** and **Supp. Fig. 1B-D**). Nuclear localization of α-syn Ser129P was also reported by a recent study using phospho-specific antibodies (Koss et al., 2022), and this smaller phosphorylated pool may be difficult to appreciate with pan-α-syn antibodies that invariably highlight the much more abundant presynaptic population. Regardless, the restricted pattern of brain α-syn Ser129P staining suggests a tight posttranslational regulation. Since levels of α-syn at synapses is linked to its function (Burre et al., 2018; Larsen et al., 2006; Nemani et al., 2010; Scott et al., 2010), we first asked if presynaptic abundance of α-syn is altered by Ser129P. Towards this, we transiently transfected GFP-tagged h-α-syn in cultured hippocampal neurons from α-syn null mice – to eliminate potential confounding effects of endogenous mouse α-syn – and used a ratio-metric imaging paradigm that accounts for variabilities in transgene expression and robustly reports presynaptic accumulation (Gitler et al., 2004) (**Fig. 1C-D**). Serine phosphorylation of α-syn occurs at two sites – Ser-87 and Ser-129 (**Fig. 1E**), although only the 129-site is predominantly phosphorylated in PD. Sequence alignment indicates that while Ser-87 is variably conserved across species, Ser-129 is highly conserved in mammals (**Supp. Fig. 1E**).

**Figure 1:**
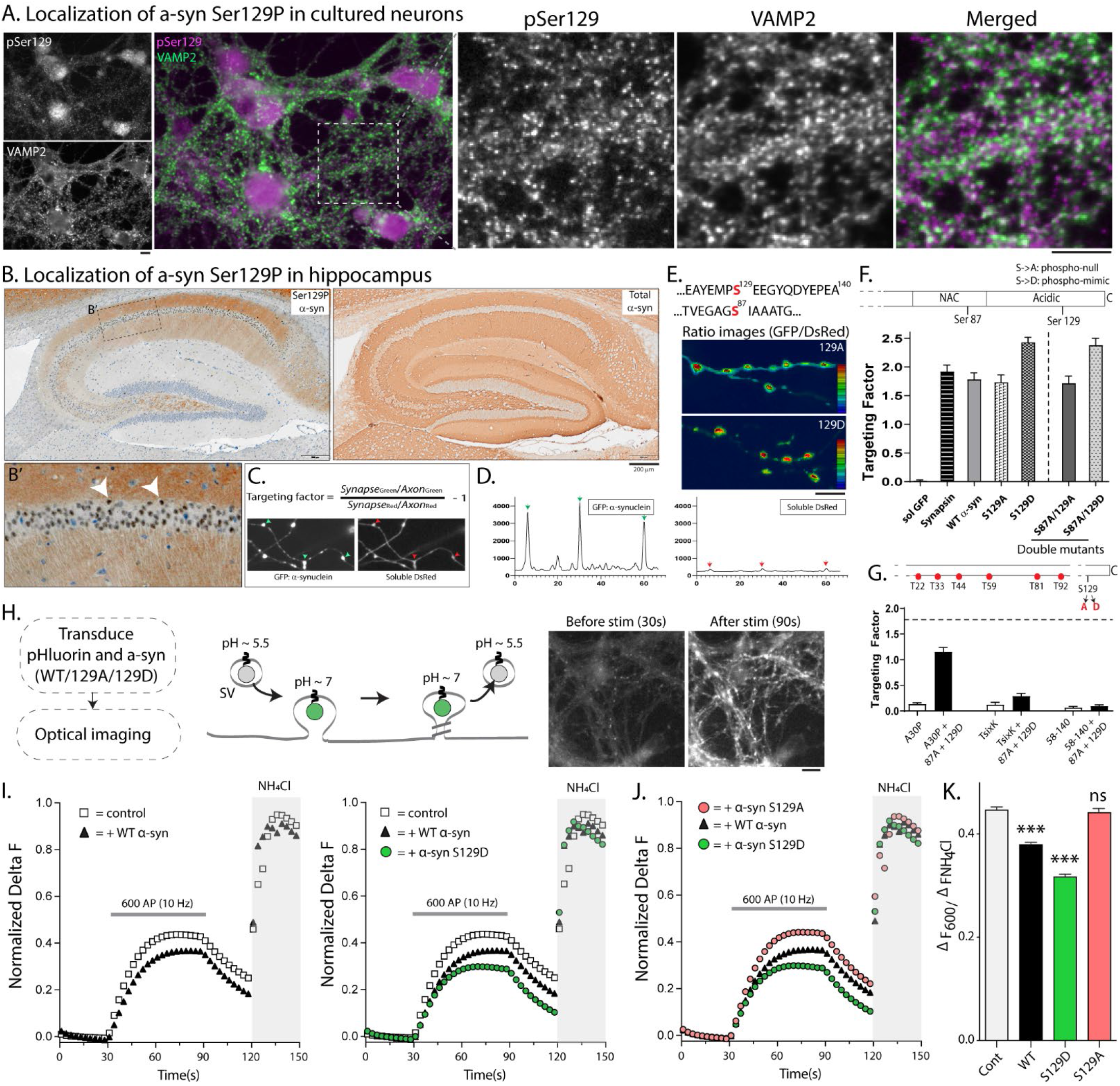
Distribution of α-syn Ser129P and its role in activity-dependent SV recycling. (A) Cultured hippocampal neurons were stained with antibodies to α-syn Ser129P and VAMP2 (to mark synapses). Note α-syn Ser129P is localized to both nuclei and synapses (also see **Supp. Fig. 1A**). Scale bars = 10 µm. (B) Mouse brain sections were stained with α-syn Ser29P and total α-syn. Note α-syn Ser129P staining in both synapses and nuclei – white arrowheads in B’ mark pSer129+ nuclei. For full characterization of the pSer129 antibody, see **Supp. Fig. 1B-D** and Methods. **(C)** <u>Top</u>: Formula for evaluating presynaptic targeting (see Gitler et al., 2004). <u>Bottom</u>: Exemplary images of neurons from α-syn null mice that were co-transfected with GFP-tagged α-syn and soluble DsRed (volume marker), showing relative enrichment of GFP/DsRed at each bouton. Linescans from these images are shown in (**D)** – note that red/green arrowheads in **(C, D)** represent same boutons. **(E)** <u>Top</u>: Serine phosphorylation sites in α-syn. <u>Bottom</u>: Representative images from synaptic targeting assays showing increased targeting of phospho-mimic (129D) α-syn. **(F)** Quantification of synaptic targeting data for phospho-incompetent (129A) and phospho-mimic (129D) mutants. While both α-syn and synapsin (another cytosolic presynaptic protein) localized to synapses, targeting was substantially increased by mimicking phosphorylation at the 129-site, and this was independent of 87A phosphorylation status (∼ 500-1000 boutons from 6-15 coverslips were analyzed for each condition; **p<0.001). **(G)** Decreased presynaptic targeting of the familial PD mutant A30P and TsixK (a synthetic mutant that disrupts membrane-binding, see text) was augmented by the 129D (but not 129A) mutant, and as expected, the N-terminus was required for targeting (∼ 600-900 boutons from 8-20 coverslips were analyzed for each condition; ***p<0.0001). **(H)** Schematic of pHluorin assay to evaluate SV-recycling (left); representative pre/post-stimulation images from vGLUT:pHl in cultured mouse hippocampal neurons transduced with vGLUT:pHl (right, see methods for details). Scale bar = 10 µm. **(I)** Fluorescence fluctuations of vGLUT:pHl in presence of WT and phospho-mutant α-syn. WT-α-syn attenuates SV-recycling in pHluorin assays as previously reported, and Ser129 phospho-mimic mutant (S129D) further augments this suppression. However, preventing α-syn Ser129P (S129A) completely abrogates the α-syn-induced synaptic attenuation **(J)**. **(K)** Quantification of the recycling pool size in all pHluorin experiments (data from 12-15 coverslips, 30-50 boutons per coverslips and 4 independent cultures for each condition; ***p<0.0001 one-way ANOVA followed by Dunnett’s post hoc test; ns = not significant).

To test the effects of phosphorylation on presynaptic targeting, we transfected neurons with h-α-syn phospho-mutants where Ser-129 was either mutated to Alanine (S129A), which cannot be phosphorylated, or Aspartic acid (S129D), a residue that mimics phosphorylation and has been used in previous studies to simulate α-syn phosphorylation (McFarland et al., 2009; Weston et al., 2021). While presynaptic targeting of phospho-incompetent S129A resembled wild-type (WT) h-α-syn, targeting of the phospho-mimic (S129D) h-α-syn was substantially increased (**Fig. 1E**, left). This augmented enrichment was specifically due to phosphorylation at the Ser129 site (see “double-mutants”, **Fig. 1F**), suggesting that phosphorylation at the 129-site may act as a switch that promotes presynaptic α-syn abundance. In further support of this, mimicking Ser-129P even augmented the synaptic targeting of A30P and TsixK mutants that normally show diminished synaptic targeting (Fortin et al., 2004; Perrin et al., 2000) (**Fig. 1G**). Expectedly, Ser-129P did not augment targeting of the deletion-mutant lacking the N-terminus (residues 58-140, **Fig. 1G**), since membrane binding via the N-terminus is essential for presynaptic targeting (Burre et al., 2012; Wang et al., 2014). Collectively, the data suggest that though Ser129P is not required for presynaptic targeting of α-syn (since phospho-incompetent mutants target normally) dynamic Ser129 phosphorylation and dephosphorylation may regulate the association of α-syn with synaptic membranes.

Modest over-expression of h-α-syn suppresses activity-dependent SV-recycling in optical pHluorin-assays that report the synaptic exo/endocytic cycle (Atias et al., 2019; Nemani et al.; Sun et al., 2019; Wang et al., 2014), leading to the concept that α-syn is a physiologic attenuator of activity-induced neurotransmitter release. Suppression of neurotransmitter release is also an early feature of PD models (Lundblad et al., 2012), suggesting that understanding the physiologic role of α-syn may be relevant for pathology. We developed a lentiviral-based system for expressing vGLUT1:pHluorin and h-α-syn in cultured hippocampal neurons – where essentially all synapses expressed the transduced transgenes – and tested the effects of modest (∼ 1.5-fold) over-expression of WT and phospho-mutant h-α-syn on SV-recycling (**Fig. 1G** and **Supp. Fig. 1F**). Since h-α-syn suppresses SV-recycling in pHluorin-assays, and phospho-mimic (S129D) h-α-syn increases synaptic α-syn levels, we expected over-expressing α-syn S129D to further attenuate SV-recycling, and this was indeed the case (**Fig. 1I**). Surprisingly however, even more than two-fold over-expression of phospho-incompetent h-α-syn (Ser129A) did not have any effect on activity-induced SV-recycling (**Fig. 1J** – all pHluorin data quantified in **Fig. 1K**). Taken together, these experiments show that serine phosphorylation at the 129-site is required for α-syn-induced synaptic attenuation, implying an unexpected physiologic role for this post-translational modification.

The α-syn-induced synaptic attenuation in pHluorin assays is activity-dependent, and neuronal activity and calcium influx is known to induce phosphorylation of a number of synaptic proteins (Kohansal-Nodehi et al., 2016; Silbern et al., 2021). Thus we first asked if neuronal activity also augmented α-syn Ser129P. For these experiments, we enhanced activity in cultured hippocampal neurons using chemical (4-aminopyridine or 4-AP, a voltage-gated K^+^ channel blocker (Thesleff, 1980) or electrical stimulation (**Fig. 2A** and **Supp. Fig. 2)**, and biochemically examined α-syn Ser129P levels. As shown in **Fig. 2B**, 4-AP treatment led to a gradual increase in Ser129P α-syn levels without a change in total α-syn. Similarly, electrical activity (600AP at 10Hz – resembling pHluorin experiments) also led to a selective increase in Ser129P levels, which was blocked by the sodium channel blocker tetrodotoxin (TTX, **Fig. 2C**). Previous studies have implicated polo-like kinase 2 (PLK2) in α-syn Ser129P (Inglis et al., 2009; Mbefo et al., 2010; Weston et al., 2021), and the PLK2 inhibitor BI2536 also prevented the phosphorylation induced by electrical activity (**Fig. 2C**). Activity-induced increase in α-syn Ser129P – with no change in total α-syn levels – was also seen in vivo in mouse brains after intraperitoneal injection of 4-AP (**Fig. 2D-F**). Collectively, the above data support a model where activity-induced α-syn Ser129P is required for α-syn functions at the synapse, but molecular mechanisms are unclear. Previous studies have shown that activity-induced phosphorylation or dephosphorylation can regulate protein-protein interaction and synaptic function. One of the best studied examples is dynamin, a protein with critical roles in endocytic membrane fission. Neuronal activity induces dephosphorylation at Ser-774/778 of dynamin, which augments its association with syndapin and promotes SV endocytosis at times of elevated neuronal activity [reviewed in (Ferguson and De Camilli, 2012)]. In other examples, serine phosphorylation of the SNARE SNAP25 accelerates SV recruitment and SNARE-complex formation (Nagy et al., 2002), and phosphorylation of syntaxin at Ser14/188 regulates its interaction with Munc18-1 (Rickman and Duncan, 2010; Tian et al., 2003). Thus we asked if Ser129P affected the interaction of α-syn with its binding partners.

**Figure 2:**
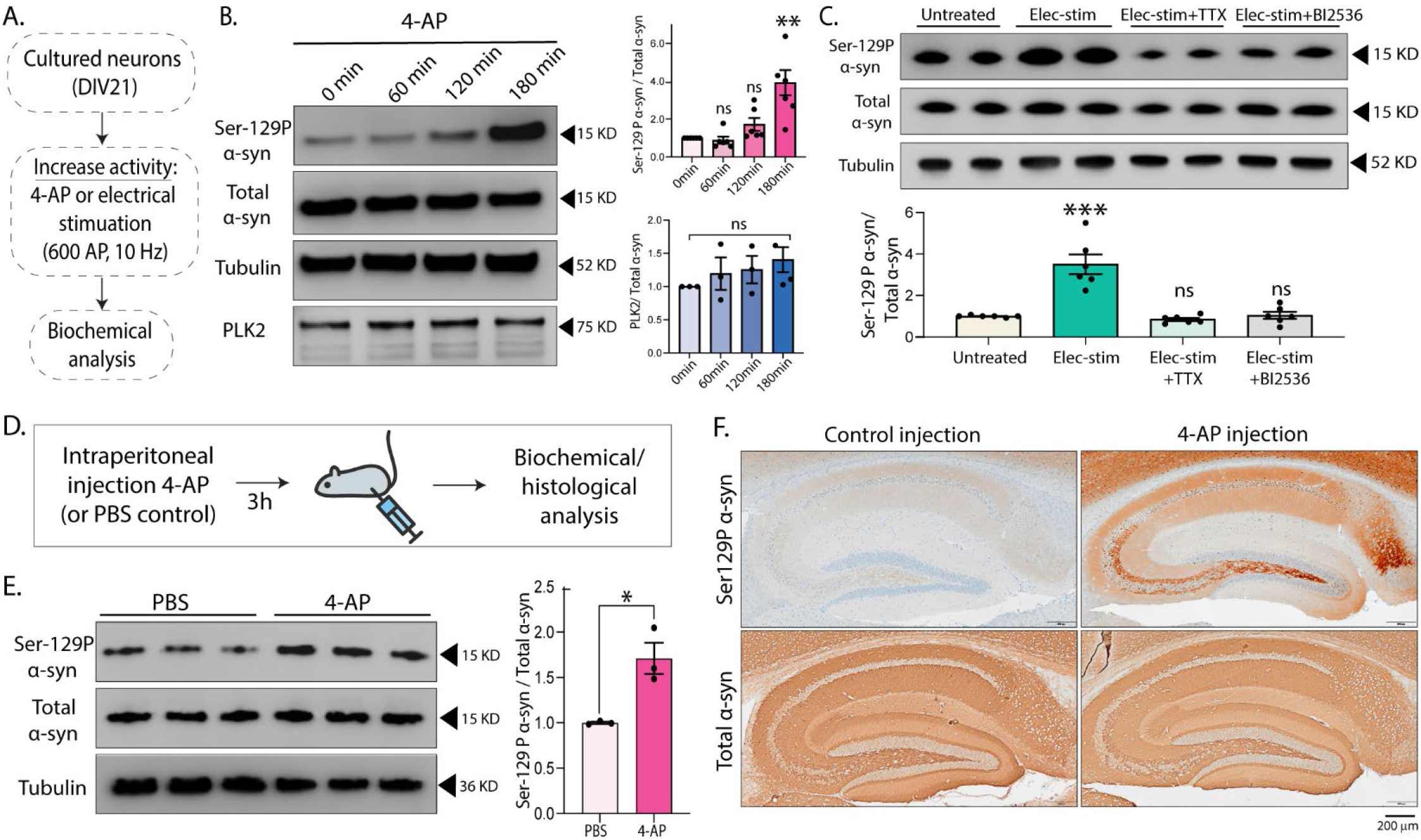
Neuronal activity augments Ser129 phosphorylation. **(A)** Experiments to induce activity in cultured neurons by chemical (4-AP, see text) or electrical (600AP) stimulation. **(B)** Western blots showing time-dependent increase in α-syn Ser129P after 4-AP treatment; quantified o right (n=6 for Ser129-P α-syn/ total α-syn and n=3 for PLK2/ total α-syn – independent experiments on different days). **(C)** <u>Top</u>: Western blots showing increased Ser129P after 600AP/10Hz stimulation (resembling vGLUT:pHl assays, see Methods). Note that pre-incubated with 1 µM TTX or 50 nM PLK2 inhibitor BI2536 (putative kinase for Ser129P) for 15min blocked Ser129P (two biological replicates shown). <u>Bottom</u>: Quantification of the western blots. **(D)** Inducing neuronal activity in vivo by intraperitoneal 4-AP injections. Mice (8 wks old) were injected with either PBS or 4-AP (2 mg/Kg dose) and sacrificed after 3h. Mouse brains were processed for western blots and histological analysis. **(E)** <u>Left</u>: Western blots from three biological replicates. <u>Right</u>: Quantification of western blots (n=3 mice). All data are means ± SEM (\**p* < 0.1, \*\**p* < 0.01, ***,*p* < 0.001 by Student’s *t* test).

VAMP2/synaptobrevin2 is a binding partner of α-syn (Burre et al., 2010), and we recently reported that binding of α-syn to VAMP2 and synapsin was essential for h-α-syn-induced synaptic attenuation in pHluorin assays (Atias et al., 2019; Sun et al., 2019). Accordingly, we wondered if serine phosphorylation at the 129-site also regulated α-syn association with VAMP2 and synapsin. Strikingly, co-immunoprecipitation (co-IP) experiments in neuro2a cells showed that interaction of h-α-syn with both VAMP2 and synapsin was essentially eliminated when the Ser129-site was rendered phospho-incompetent (**Figs 3A-C**; also see **Supp. Fig. 3A-B**).

**Figure 3:**
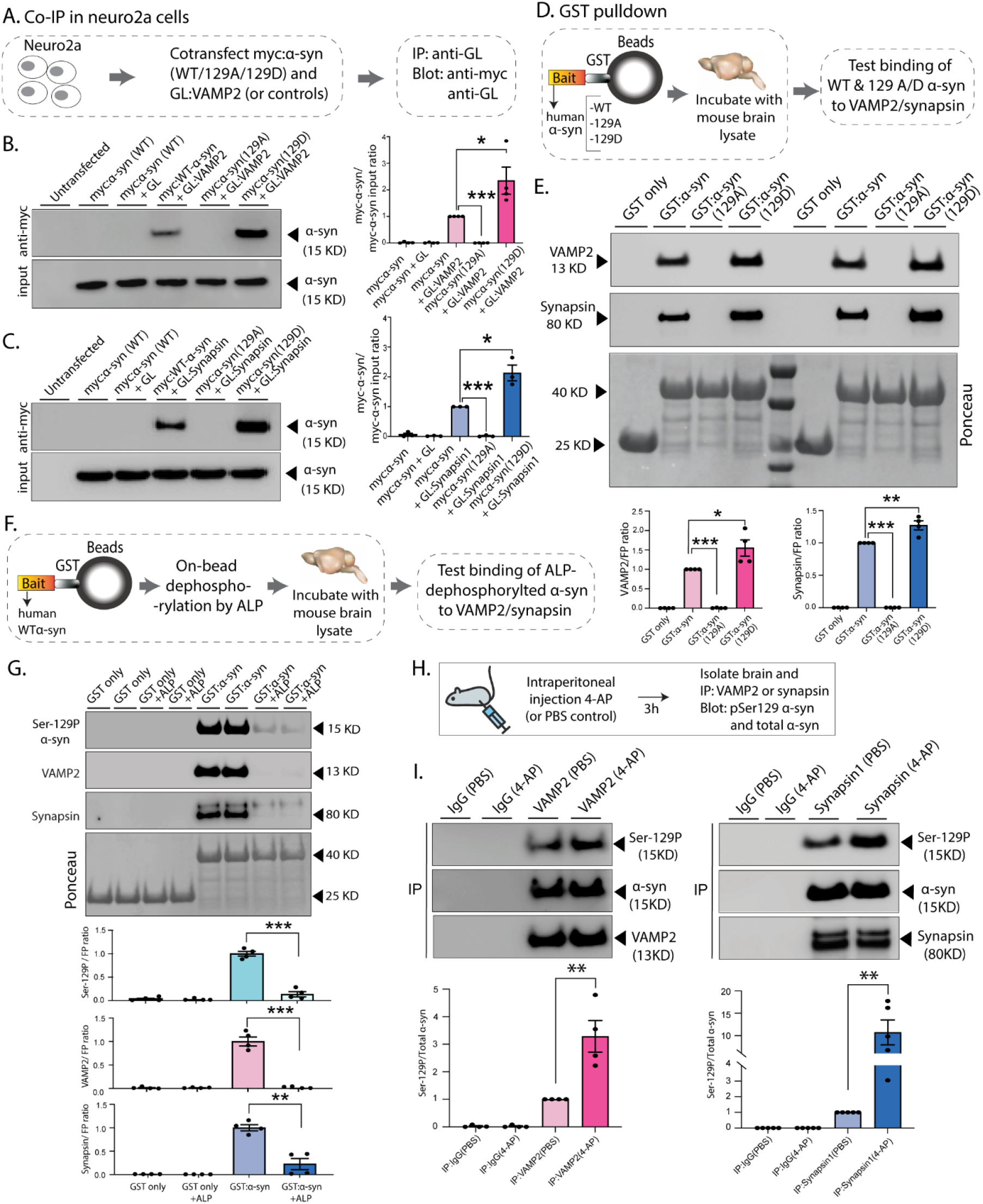
Ser129P regulates association of α-syn with functional binding partners VAMP2 and synapsin. **(A**) Workflow for co-immunoprecipitation (Co-IP) experiments in neuro2a cells. **(B, C)** Western blots from co-IP experiments show that both VAMP2 **(B)** and synapsin **(C)** was co-IP’d with WT-α-syn, but not phospho-incompetent (129A) α-syn. Mimicking Ser129P (S129D) augmented this interaction – quantified on right (n=4 for myc α-syn/GL-VAMP2 and n=3 for myc α-syn/GL-Synapsin co-IP). **(D)** Workflow for pulldown of GST-tagged WT/129A/129D α-syn after incubation with mouse brain lysates. Equivalent amounts of immobilized GST α-syn (or its phospho variants) were used. **(E)** Samples from GST-pulldown were analyzed by NuPAGE and immunoblotted with antibodies against VAMP2 (blot in top panel) and synapsin (blot in middle panel); two biological replicates are shown. Ponceau staining (bottom panel) shows equivalent loading of fusion proteins. Note that preventing Ser129P (S129A) blocked interaction with VAMP2 and synapsin. Western blots quantified below (n=4). **(F)** Workflow for on-bead α-syn dephosphorylation. Beads containing GST-tagged WT-α-syn (or S129A/D) were pretreated with alkaline phosphatase (2.5 units per µg of protein) for 3h before pulldown. **(G)** Samples from GST-pulldown were analyzed by NuPAGE and immunoblotted with antibodies against Ser-129P α-syn, VAMP2 and synapsin. Ponceau staining (bottom panel) shows equivalent loading of fusion proteins. Note that α--syn dephosphorylation attenuates VAMP2 and Synapsin interaction. Western blots quantified below (n=4). **(H)** Workflow for chemical activity induction (4-AP injections), followed by co-IP experiments to detect interactions between VAMP2/synapsin and ser-129P α-syn. **I)** Note that after 4-AP injections, a higher fraction of Ser129p α-syn was co-IP’d with VAMP2 (left) and synapsin (right). Western blots of VAMP2 or Synapsin co-IP’d with Ser129P α-syn quantified below (n=4 and n=5 respectively). All data presented as mean ± SEM (\**p* < 0.01, \*\**p* < 0.001, \*\*\**p* < 0.0001, t test).

To evaluate the interaction of phospho-incompetent (129A) and phospho-mimetic (129D) h-α-syn in the context of the brain, we incubated mouse brain lysates with beads carrying GST-tagged recombinant h-α-syn WT/S129A/S129D, and analyzed proteins bound to h-α-syn by western blotting (**Fig. 3D**). While performing these experiments, we noticed that the recombinant WT-α-syn expressed in bacteria was already phosphorylated at the 129D-site, likely by endogenous bacterial kinases (see characterization of purified proteins in **Supp. Fig. 3C-D**). Consistent with the immunoprecipitation experiments, brain VAMP2 and synapsin also failed to bind S129A (phospho-incompetent) h-α-syn in GST-pulldown assays, whereas robust binding was seen with both WT-α-syn (endogenously phosphorylated) and the S129D (phospho-mimetic) α-syn (**Fig. 3E**). Authentic on-bead dephosphorylation of h-α-syn in these experiments using alkaline phosphatase also dramatically attenuated α-syn interaction with VAMP2 and synapsin, strongly implying that this phosphorylation event is critical for protein-protein interactions (**Fig. 3F-G**). Since neuronal activity augments α-syn Ser-129P (**Fig. 2**), a prediction is that under conditions of enhanced neuronal activity, a higher fraction of phosphorylated α-syn would associate with VAMP2 and synapsin; and indeed, this was the case in mice injected with 4-AP (**Fig. 3H-I**). Taken together, the data indicate that activity dependent serine phosphorylation at the 129-site of α-syn triggers its association with the functional interacting partners VAMP2 and synapsin.

The α-syn C-terminus (residues 96-140) has emerged as an important site for protein-protein interactions (Burre et al., 2010; Sun et al., 2019), and both VAMP2 and synapsin bind to this region (**Supp. Fig. 4A**). Previous cell-free experiments with purified proteins and vesicles have shown that α-syn facilitates the association and clustering of small synaptic-like vesicles, and that VAMP2 is necessary for this clustering (Diao et al., 2013; Sun et al., 2019). Direct tethering of adjacent vesicles by α-syn has also been described (Fusco et al., 2016b; Lautenschlager et al., 2018), and α-syn-dependent binding and mobilization of SVs may be important clues to its function. Since both VAMP2 and synapsin are associated with SVs, one interpretation of our biochemical experiments is that Ser129P facilitates the association of α-syn with SVs enriched in VAMP2/synapsin. While bulk biochemical approaches have estimated average protein-numbers on SVs (Takamori et al., 2006), quantitative measurements at a single-SV level indicate that the protein composition of individual SVs is variable, with VAMP2 levels showing significant inter-vesicle variability (Mutch et al., 2011). Other studies are also consistent with a variance in protein numbers between SVs (Gracz et al., 1988; Hayashi et al., 2008), which likely occurs during membrane retrieval and possibly other sorting mechanisms that remain poorly defined.

Accordingly, we first asked if phosphorylation at the Ser129-site favored association of α-syn with SVs enriched in VAMP2 and synapsin. For these experiments, we incubated GST-tagged WT/129A/129D h-α-syn with SV-fractions isolated from mouse brains (**Fig. 4A**, see protocol in **Supp. Fig. 4B**). Note that synucleins are absent from these purified SV-fractions as described previously (Kahle et al., 2000) – presumably due to relatively weaker association with SVs – eliminating potential confounding effects of mouse α-syn in these experiments (**Fig. 4B**). Indeed, VAMP2/synapsin enriched SVs preferentially bound to WT (natively phosphorylated) and 129D (phospho-mimic) h-α-syn, compared to phospho-incompetent (S129A) h-α-syn, while protein-levels of SV2 – an integral membrane protein that is equally distributed amongst SVs (Mutch et al., 2011) – was similar in all three conditions (**Fig. 4C**). Though Ser129P led to a clear increase in VAMP2/synapsin binding, the relative affinities of synapsin and VAMP2 containing SVs to phosphorylated and dephosphorylated α-syn was variable, and we reasoned that this might be due to the additive influence of the N-terminus of α-syn, which is also expected to bind SVs in this assay.

**Figure 4:**
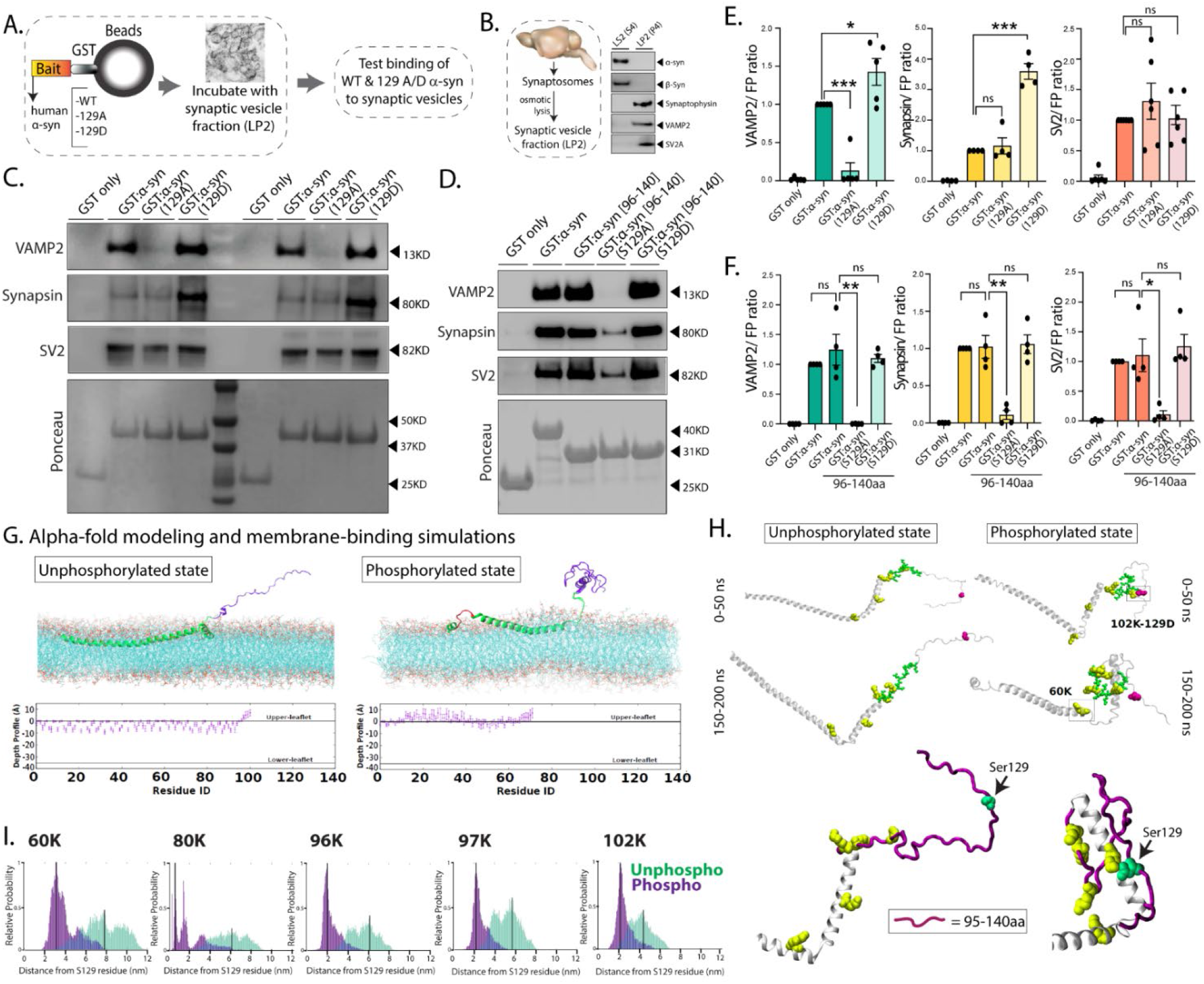
Ser129P regulates association of α-syn with SVs enriched in VAMP2 and synapsin. **(A)** Workflow for pulldown of GST-tagged WT/129A/129D α-syn after incubation with SV-fractions (LP2) from mouse brain lysates. Equivalent amounts of immobilized GST α-syn (or its phospho variants) were used. **(B)** <u>Left</u>: Overall protocol for isolating synaptosomes, see Supp. Fig. 4B for details. <u>Right</u>: Western blots from the pellet (LP2, 15 µg) and supernatant (S4) were analyzed by western blots. Note that mouse α-syn is not present in these enriched fractions, eliminating potential confounding factors. **(C, D)** Samples from GST-pulldown [GST-tagged WT-α-syn in **(C)** and GST-tagged 96-140-α-syn – binding region of VAMP2/synapsin – in **(D)**] were analyzed by NuPAGE and immunoblotted with antibodies against VAMP2 (blot in top panel), synapsin (blot in middle panel), and a housekeeping protein SV2 (bottom panel). Ponceau staining (bottom panel) shows equivalent loading of fusion proteins. Two biological replicates are shown in **(C)**. Note that in all cases, preventing Ser129P blocks the association of α-syn with VAMP2/synapsin enriched SVs, and residues within 96-140 are sufficient to mediate this effect (see text for more details). **(E)** Quantification of blots in **(C)**: VAMP2 (n=5), synapsin (n=4), and SV2 (n=6). **(F)** Quantification of blots in **(D)**: VAMP2, synapsin, and SV2 (n=4). All data presented as mean ± SEM (\**p* < 0.1, \*\**p* < 0.01, \*\*\**p* < 0.001, *t* test). **(G)** Structures of α-syn were generated using Alphafold2-drived Colabfold, and membrane-binding was simulated (see text and methods). <u>Top</u>: Near-final snapshots from simulations showing depth-profiles of membrane embedded unphosphorylated (left) phosphorylated (right) states of α-syn. <u>Bottom</u>: Quantitative residue-wise depth-profile (mean +/-SEM of each residue) shows membrane-embedding in the two states. Note that the C-terminus appears folded in the phosphorylated state. **(H)** Representative snapshots from early (0-50 ns) and late (150-200 ns) stages of the simulation show that in the phosphorylated state, five positively charged lysine residues within the 95-140 residues (60K, 80K, 96K, 97K and 102K – marked in yellow) crowd around the Ser129-site (marked by an arrow in the zoomed images below). The distances between these five lysine residues and the Ser129-site across the simulations is quantified in **(I)**. Note the greater proximity of the five lysines to Ser129 in the phosphorylated state.

In support of this, similar experiments using only the VAMP2/synapsin binding region of α-syn (residues 96-140) showed an almost binary effect of Ser-129-P on binding to VAMP2/synapsin enriched SVs (**Fig. 4D**), confirming that this phosphorylation event is required for such protein-protein interaction. Note that in this scenario, SV2 levels are expectedly low in the 129A condition because *only* SVs containing VAMP2/synapsin would bind to the α-syn C-terminus, and these vesicles are also expected to have SV2 which is equally distributed amongst SV (Mutch et al., 2011) (gels from **Figs 4C-D** quantified in **Figs. 4E-F**). Taken together, the data suggest that phosphorylation and dephosphorylation at the Ser-129 site of α-syn regulates its association with a subset of SVs enriched in VAMP2 and synapsin, and that this effect is mediated by the C-terminus of α-syn (residues 96-140).

How does a single phosphorylation-event modulate α-syn interactions? Since the Ser129P-induced differential binding of α-syn with its interacting partners is solely mediated by residues 96-140 (**Fig. 4D**), one possibility is that α-syn Ser129P induces conformational changes within the C-terminus that stabilize this region, facilitating binding to interacting partners. Indeed, there are many examples in biology where phosphorylation alters protein conformation and protein-protein binding (Johnson and Lewis, 2001). To explore the effects of Ser129P on α-syn structure, we used AlphaFold2-based modeling and membrane-binding simulations. AlphaFold2 uses machine learning algorithms that incorporate both bioinformatics and physical knowledge of protein structures – leveraging multi-sequence alignments – to accurately predict three-dimensional structures at atomic resolution; and has emerged as a useful tool in predicting three-dimensional structures of proteins (Jumper et al., 2021). To model putative changes in α-syn structure upon Ser129P, we used ColabFold, a free and accessible platform based on AlphaFold2 and RoseTTAFold, which reproduces and implements many of the ideas incorporated by AlphaFold2 (Mirdita et al., 2022). Since regulation of α-syn Ser129P is dynamic (Okochi et al., 2000; Weston et al., 2021), we considered two states in our simulations – one where the Ser129 remains unphosphorylated, and another where h-α-syn is constitutively phosphorylated (phospho-mimic S129D); employing models that had the highest confidence values in ColabFold (**Supp. Fig. 4C-D**). Finally, we set up an atomic simulation where resultant ColabFold structures were allowed to associate with artificial bilayers resembling SV membranes under physiologic conditions, and penetration of h-α-syn into the bilayers was determined (see **Supp. Fig. 4E-F** and methods). For each scenario, only the near-final stabilized, membrane-embedded conformation from the simulations were considered (the last 20ns, ∼10% of the simulation, averaging 10^7^ time-points at 2 femto-second resolution – see methods).

Though there were changes in membrane-binding in the phosphorylated state, the N-terminus was largely embedded into the membrane, and there was little change in the overall conformation of the vesicle-binding surface (**Fig. 4G** and **Supp. Fig. 4D**). Moreover, since membrane-binding of α-syn is thought to mainly rely on the first ∼ 20 residues (Fusco et al., 2016a), these changes are not expected to have a significant effect on SV-binding. However, the simulations revealed that the negative charges added to amino acid side chains by Ser129P facilitated interactions of this site with five positively charged lysine residues within the VAMP2/synapsin-binding region (60K, 80K, 96K, 97K, and 102K) – see snapshots from simulation in **Fig. 4H** and folding-kinetics in **Supp. Movie 1**. Interestingly, these interactions tend to crowd the five lysines around the 129-site (see zoomed exampled from simulation, **Fig. 4H** – bottom panel), and we speculate that such associations might stabilize the binding region of VAMP2 and synapsin (96-140 residues) and facilitate protein-protein interactions. Quantitative analyses of inter-residue distances highlight the proximity of C-terminus lysine residues to the 129-site during the simulation when α-syn is in the phosphorylated state (**Fig. 4I**). Taken together, our data from cellular, in vivo, cell-free systems, and molecular simulations strongly suggest that the pathology-associated Ser129 modification has a physiologic role at the synapse, which has broad implications for α-syn pathophysiology and drug development.

## Methods

### Animals, reagents and DNA constructs

All animal studies were performed in accordance with University of California guidelines. Primary hippocampal cultures were obtained from timed-pregnant female CD-1 mice (Charles River); biochemical and in vivo experiments were done using C57BL/6J mice (Jackson Labs). The following antibodies were used for immunofluorescence experiments: Ser129P α-syn (Abcam #ab51253, 1:100), NeuN (Abcam #ab279297, 1:1000), VAMP2 (Synaptic systems #104211, 1:1000). The following antibodies were used for biochemistry experiments: Ser129P α-syn (Abcam #ab51253, 1:800), total α-syn (BD Bioscience #610787, 1:400), tubulin (Abcam #ab7291,1:10000), PLK2 (Abcam #ab137539,1:10000), synaptophysin-1 (Synaptic systems #101011,1:500), VAMP2 (Synaptic systems #104211, 1:500), SV2A (Abcam #ab32942,1:500), synapsin-1 (Abcam #ab254349, Synaptic systems #106104), c-myc (Sigma #M4439, 1:500), GFP (Abcam#ab290, 1:5000). Chemicals used: 4-Aminopyridine (4-AP, Sigma #A78403), tetrodotoxin (TTX) (Cayman#14964), 6-cyano-7-nitroquinoxaline-2,3-dione (CNQX, TOCRIS bioscience #0190), D,L-2-amino-5-phosphonovaleric acid (AP5, TOCRIS bioscience #0105), and BI2536 (Selleckchem #S1109). Antibodies used for tissue immunohistochemistry are described in a separate section. For tagged proteins, synthetic DNA blocks (IDT) or PCR products coding for human WT α-syn, α-syn S129A, α-syn S129D, VAMP2, and synapsin1 were subcloned into the pCCL-mScarlett, pCCL-Green Lantern, or pGEX-KG vectors. Synthetic DNA blocks (IDT) coding for the truncated α-syn variants were subcloned into the pGEX-KG vector. All cloning was performed using Gibson cloning (NEB). Constructs obtained from other laboratories are noted in the acknowledgements. All constructs were verified by sequencing.

### Hippocampal Cultures, lentivirus production, vGLUT1-pHluorin imaging and analysis

Primary hippocampal cultures were obtained from P0-P1 mouse pups using standard procedures as described previously (Roy et al., 2012; Scott and Roy, 2012). For synaptic targeting experiments, neurons were transiently transfected on DIV-13 using Lipofectamine 2000 (Invitrogen) and imaged at DIV 17-21 as described previously (Wang et al., 2014). For lentivirus production, HEK293T cells were maintained in DMEM+Glutamax supplemented with 10% FBS and 1% penicillin-streptomycin. 6Ð10^6^ cells were plated in 15cm dishes (Genessee Scientific) 14-16 hrs. before transfection. HEK cells were transfected with the targeting plasmid and two helper plasmids (psPAX2 and pMD2.G) at a 2:1.5:1 molar ratio using ProFection (Promega). Two to three days later, the supernatant was collected and concentrated using LentiX (Takara). The viral pellet was resuspended in 1/100th supernatant volume of sterile HBSS and stored at –80°C. For viral transductions, lentiviruses were added to each well of neurons at DIV 3 at MOI=5. In all cases, there was almost 100% transduction with the lentiviruses, as confirmed by

For vGLUT1:pHluorin experiments, cultured neurons were plated at a density of 60,000 cells/cm^2^. DIV-3 neurons were infected with lentiviruses carrying α-syn tagged at the C-terminus to mScarlett, or mScarlett alone (multiplicity of infection or MOI=2.5). Subsequently, lentiviruses carrying vGLUT1:pHluorin (MOI=2.5) were added on DIV-5 and the transduced neurons cultured to maturity (DIV17-DIV21) before imaging. 100% infection efficiency of the mScarlett tagged constructs was confirmed before using the coverslip for pHluorin experiments. Neurons were imaged live using an inverted motorized epifluorescence microscope (Olympus, IX81) fitted with a Prime 95B camera. The coverslips were mounted into a Chamlide EC magnetic chamber (Live cell Instrument, ON, Canada) in Tyrode solution (pH 7.4) containing (in mM): 119 NaCl, 2.5 KCl, 2 CaCl_2_, 2 MgCl_2_, 25 HEPES, 30 glucose, 10 μM 6-cyano-7-nitroquinoxaline-2,3-dione (CNQX, TOCRIS bioscience #0190), and 50 μM D,L-2-amino-5-phosphonovaleric acid (AP5, TOCRIS bioscience #0105). NH_4_Cl perfusions were done with 50 mM NH_4_Cl in substitution of 50 mM NaCl (pH 7.4). For field-stimulation, 10 V/cm pulses were applied at 10Hz for 60 seconds using a Model 4100 Isolated High Power Stimulator (A-M systems, Sequim, WA). Incident excitation (Lumencor LED, Spectra X) was attenuated 10-fold, and images were acquired with 500 ms exposures at three second intervals for three minutes. For analysis, regions of interest (ROIs) were placed on each bouton and average intensities were obtained for each frame within the time-lapse. F_max_ was defined as the maximal fluorescence after NH_4_Cl perfusion. Baseline F_0_ was defined as the average fluorescence of the initial 10 frames before stimulation [(F_0_ = average (F_1_:F_10_)]. Fluorescence intensity of a bouton at a given time point (F) was normalized to F_0_ and F_max_ and expressed as (F-F_0_) / (F_max_-F_0_).

### Immunofluorescence and immunohistochemistry

For immunostaining, DIV17-21 neurons were fixed in 100% ice-cold methanol solution for 15 min at room temperature, followed by extraction in 0.1% Triton X-100 for 10 min, blocking in 10% BSA for 1 h at room temperature, and overnight incubation at 4°C with primary antibodies. After washing with 1x PBS, neurons were blocked again for 30 min at room temperature and incubated with secondary antibodies (Alexa Fluor antibodies from Invitrogen, 1:500) for 1 hour at room temperature. Images were acquired at 40X. Z-stack images were obtained as previously described (Scott and Roy, 2012) and all images were acquired and processed using MetaMorph software. For immunohistochemistry, 5 µm tissue sections were stained with antibodies to total α-syn (1:15,000, Synaptic Systems) and Ser129P α-syn (Cell Signaling, 1:3000, Cat #23706). Slides were stained on a Ventana Discovery Ultra (Ventana Medical Systems, Tucson, AZ, USA). Antigen retrieval was performed using CC1 (Tris-EDTA based; pH 8.6) for 40 minutes at 95°C. The primary antibodies were incubated in the sections for 32 minutes at 37°C. To confirm specificity of the Ser129P α-syn antibody, the epitope was dephosphorylated using lambda protein phosphatase (P0753; New England Biolab; Ipswich, MA) for 2 hours at 37°C according to the manufacturer’s protocol (adapted for use on the Ventana Discovery Ultra). Primary antibodies were detected using the OmniMap systems (Ventana Medical systems). Antibodies were visualized using DAB as a chromogen, followed by hematoxylin as a counterstain. Slides were rinsed, dehydrated through alcohol and xylene, and mounted on coverslips.

### Biochemical assays and evaluation

#### Preparation of Brain and Neuro2A Lysates

Whole mouse brains were homogenized with a Dounce tissue grinder in neuronal protein extraction reagent (N-PER) (ThermoFisher) containing protease/phosphatase inhibitors (Cell Signaling #5872). Triton X-100 was added to a final concentration of 1%, and the samples were incubated with rotation for 1 hour at 4 °C. Samples were centrifuged at 10,000 × *g* for 10 min at 4 °C, and the supernatant was collected. To obtain Neuro2A lysates, cells were washed with 1X PBS 3 times and incubated 5min on ice in the presence of N-PER reagent supplemented with protease inhibitors. Samples were centrifuged at 10,000 × *g* for 10 min at 4 °C to remove cellular debris. After obtaining the brain and Neuro2A lysates we measure protein concentration (DC™ Protein Assay Kit II, Biorad) and samples were used in subsequent experiments.

#### Preparation of an Enriched Synaptic Vesicle Fraction

Mouse brain tissue was homogenized in a buffer containing 0.32M sucrose in PBS and supplemented with protease/phosphatase inhibitors. The homogenate was centrifuged at 1,000 × *g* for 10 min at 4 °C to remove nuclei and cellular debris (P1). The supernatant (S1) was centrifuged at 15,000 × *g* for 15 min at 4 °C to yield a crude synaptosomal pellet (P2). This pellet was hypo-osmotically lysed in water containing protease inhibitors for 5 min at 4 °C and passed through both a 22- and 27½-gauge needle (10 times each). This suspension was centrifuged at 23,000 × *g* for 22 min at 4 °C, and the resulting supernatant LS1(S3) was centrifuged again at 174,900 × *g* for 2hrs. at 4 °C in a STi32 Beckman rotor. The final pellet called LP2(P4) contained an enriched fraction of synaptic vesicles and was used for subsequent experiments.

#### Immunoprecipitations and Western blots analysis

Immunoprecipitations were performed using 1-2 mg of total protein. Samples were incubated overnight with the indicated antibody at 4 °C, followed by the addition of 50 μl of protein G-agarose beads (ThermoFisher). Immunoprecipitated proteins were recovered by centrifugation at 2,500× rpm for 2 min, washed three times with a buffer containing PBS and 0.15% Triton X-100. The resulting pellets were resuspended in 20 μl of 1X NuPAGE LDS sample buffer (ThermoFisher #NP007) and incubated at 95 °C for 10 min. Samples were separated by NuPAGE 4 to 12% Bis-Tris polyacrylamide gels (ThermoFisher #NP0335BOX), and transferred to a 0.2 µM PVDF membrane (ThermoFisher #LC2002), using the Mini Blot Module system (ThermoFisher). PVDF membranes were first fixed with 0.2% PFA 1x PBS per 30min at room temperature. Then, membranes were washed three times for 10 min in PBS with 0.1% Tween 20 Detergent (TBST) and blocked for 1 h in TBST buffer containing 5% dry milk, and then incubated with the indicated primary antibody for 1 h in blocking solution, washed three times for 10 min each, and incubated with an HRP-conjugated secondary antibody. After antibody incubations, membranes were again washed three times with TTBS buffer and protein bands were visualized using the ChemiDoc Imaging System (BioRad) and quantified with Image Lab software (BioRad).

#### GST Pull-down Assays

Constructs containing glutathione *S*-transferase (GST) fusion proteins were expressed in bacteria, and purified as described previously (Carneiro et al., 2002). Briefly, recombinant constructs were transformed in BL21 *Escherichia coli* and treated with 0.05 mM IPTG (isopropyl-β-d-thiogalactopyranoside), and, either incubated at 37°C for 2 hours, or at room temperature for 6 hours, with shaking. The cells were harvested by centrifugation at 4500× *g* at 4 °C for 20 min and pellets were stored at -80 °C until use. For protein purification, protein pellets were resuspended in 30 ml Lysis Buffer (PBS, 0.5 mg/ml lysozyme, 1 mM PMSF, DNase, and EDTA-free protease cocktail inhibitor) for 20 min at room temperature, briefly sonicated, and removed the insoluble material by centrifugation at 20,000g at 4 °C for 30 min. The clarified lysate was incubated with 500 µl of glutathione-Sepharose 4B (AP Biotech), preequilibrated with PBS containing 0.1% Tween 20 and 5% glycerol (binding buffer), on a tumbler at 4 °C overnight. The GST-bound proteins were washed four times with 30 ml binding buffer and maintained at 4 °C. To pull down proteins from brain lysates and LP2(P4) fractions, 1-2 mg of the sample was incubated with 25-50 μg of glutathione beads containing GST fusion proteins for 12-16 hrs. The mixtures were washed three times with PBS with 0.15% Triton X-100, and then resuspended in 20 μl of 1X NuPAGE LDS sample buffer. Finally, samples were analyzed by NuPAGE and immunoblotted with the indicated antibodies.

### Multi-electrode array (MEA)

One day before plating, 6-well MEA plates (Axion Biosystems, M384-tMEA-6W) were coated with poly-D-lysine (Millipore Sigma P6407-5MG) in borate buffer, then washed six times with sterile water and aspirated prior use. 150,000 cells in a 15 µl drop were plated on a 6-well MEA plate. Cultures were maintained for 17-21 days at 37 °C in 5% CO2. On the day of recording, MEA plates were equilibrated for 10 min on the pre-warmer reader at 37 °C in 5% CO2. For spontaneous neuronal activity, we collected 2 min of spontaneous firing activity at a 12.5 kHz sampling rate and filtered with a bandpass filter from 200 Hz to 2.5 kHz. For electrically evoked extracellular activity, we activated 4 electrodes using 1000 µs stimulus duration, a voltage of 1250 mV, and a max current of 100 µA. Pulses were applied at 10 Hz for 60 seconds (600AP). Each MEA well contained 64 electrodes that detected changes in extracellular field potentials reflecting the spiking activity of neurons. Spike detection was performed with the adaptive threshold crossing algorithm with a threshold of 6× standard deviation using the AxIS software version 2.4 (Axion Biosystems, Atlanta, GA, USA). All spike trains were exported from AxIS software.

### ColabFold Protein Structure Prediction and Membrane Interaction Modeling

Structures of WT and phospho-mutant α-syn were predicted using ColabFold, which uses MMseqs2 for homology detection and multiple sequence alignment and AlphaFold2 for simulated protein folding (Jumper et al., 2021). The full-length human α-synuclein isoform 1 sequence (Uniprot, P37840-1) was input into ColabFold with 3 recycles, and the model with highest overall IDDT (Mariani et al., 2013) was chosen as representative. For modeling of the phospho-mimetic mutant of α-syn, Ser129 was replaced with an aspartate residue before input. To model the interaction between α-syn and SV-membranes, an all-atom simulation was set up, which included the previously selected ColabFold protein models and a phospholipid bilayer made up of 1-palmitoyl-2-oleoyl-sn-glycero-3-[phospho-rac-(1-glycerol)] (POPG) (Jao et al., 2008) spread in the x-y plane. The protein models were introduced as close to the membrane as possible, while ensuring no disruption of the bilayer, and then the simulation box was solvated with TIP3P water molecules, and charge neutralized by the required number of K^+^ and Cl^-^ions. Simulations were conducted using GROMACS v.2021.3 [doi: 10.5281/zenodo.5053201] with a CHARMM36 force field (Huang and MacKerell, 2013) at a temperature of 310.15K. Particle size of the simulations was 618946 and 324399 for wild-type and S129D α-synuclein, respectively. After simulation setup, energy minimization was performed using the steepest descent minimization algorithm, and the LINCS algorithm was used to achieve the constraints for hydrogen bonds (Hess et al., 1988). Equilibration of the system was performed with two steps of NVT with Berendsen temperature coupling used for temperature correction with a time step of 1 femtosecond, followed by four steps of NPT with Berendsen temperature and pressure coupling with a time-step of 1 femtosecond for the first NPT step and 2 femtoseconds for the following three. All equilibration steps were run for 125 picoseconds each. Finally, the molecular dynamics production run was performed, without any constraints, for approximately 200 ns using Nosé-Hoover temperature coupling (Evans and Holian, 1985) and Parrinello-Rahman pressure coupling (Parrinello et al., 1981), with a time-step of 2 femtoseconds. In the final model, the helicity of each residue is represented as a fraction of time spent either as a component of an α-helical segment of the protein, a 1, or otherwise, a 0. The depth profile of each protein model is represented as the position of each residue with respect to the phosphate head group, and is shown as the mean and standard deviation of the last 20 nanoseconds of the production run.

### Multiple Sequence Alignment

Protein sequences were retrieved from Uniprot or NCBI via Jalview’s sequence fetcher (Waterhouse et al., 2009), and aligned with the Clustal Omega Webserver with default parameters (Higgins and Sharp, 1988). For the a-synuclein alignment shown in Fig. X, human (uniprot OP37840), chimpanzee (uniprot P61145), baboons (uniprot A0A2I3N0Z9), marmoset (uniprot F7GY62), rat (uniprot P37377), mouse (uniprot O55042), rabbit (uniprot G1U0V2), cow (uniprot Q3T0G8), bat (uniprot A0A6J2L5S3), shrew (uniprot J7H0X3), armadillo (NCBI XP_004480597), elephant (uniprot G3T7Z3), tenrec (NCBI XP_004703317.1),chicken (uniprot Q9I9H1), zebra finch (uniprot Q4JHT6), frog (uniprot Q7SZ02), and fish (uniprot A0A090D865) sequences were used.

### Statistical Analysis

Statistical analyses were performed using Prism software (GraphPad, La Jolla, CA). Nonparametric Student’s t-test was used for comparing two groups, and one-way ANOVA for multiple groups. Results are expressed as mean±SEM. A p-value <0.05 was considered significant.

## Acknowledgements

We thank Drs. Julia George (Univ. of London), George Augustine (Nanyang Technological University, Singapore), and Shigeki Watanabe (Johns Hopkins) for the TsixK, GFP:synapsin-Ia and lentiviral VGLUT1-pHuorin constructs respectively; and Dr. Jaqueline Burre (Weill-Cornell) for the myc-tagged constructs. This work was supported by grants to Subhojit Roy from the NINDS (R01NS111978), the Farmer Family Foundation, the ASAP (Aligning Science Across Parkinson’s) network, and an NINDS P30NS047101 grant to the UCSD microscopy core. Leonardo Parra was supported by a postdoctoral fellowship from the American Parkinson’s Disease Association (APDA).

---

**Supplementary Fig. 1:**
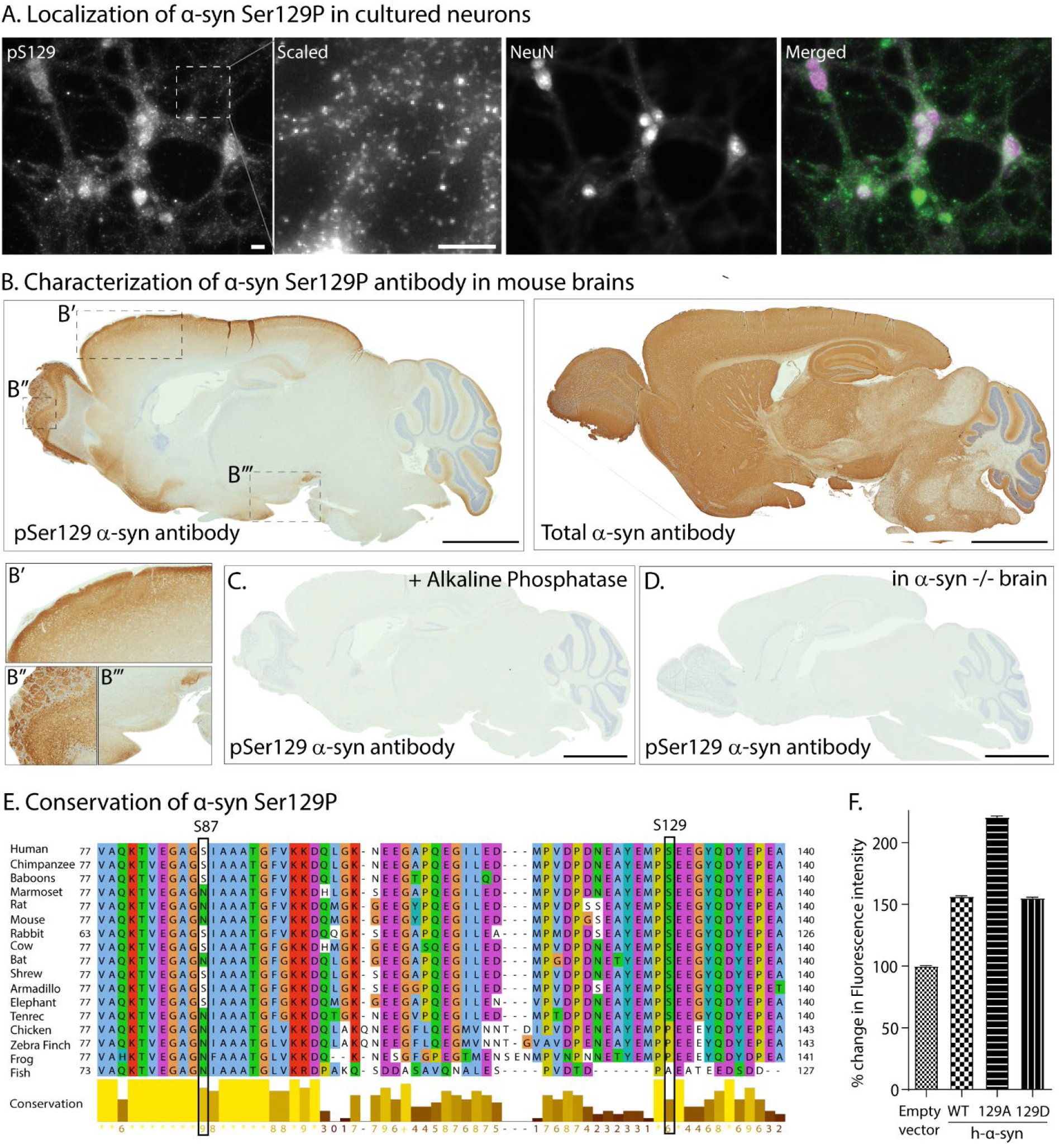
Ser129P α-syn subcellular localization, antibody characterization, evolutionary conservation, and over-expression levels of WT and phospho-mutant α-syn in vGLUT-pHl experiments. **(A)** Representative images from cultured hippocampal neurons stained with antibodies to Ser129P α-syn and NeuN (to stain neuronal nuclei). Note synaptic-like staining of Ser129P α-syn in zoomed inset (scaled to maximize bit-depth and highlight puncta), as well as nuclear staining – colocalized with NeuN in merged image. **(B)** Adjacent sections from a mouse brain were stained with antibodies recognizing Ser129P α-syn (left) and all endogenous α-syn (right). Note that unlike the staining of total α-syn – that broadly labels synapses throughout the brain – Ser129P staining is restricted to the superficial cortex, hippocampus, olfactory bulb, and dopaminergic regions. Zoomed insets **(B’, B’’, B’’’)** highlight both synaptic and nuclear staining (also see **Fig. 1B**). Scale bar = 2 mm. **(C, D)** Pre-treatment of the tissue with phosphatases eliminated the Ser129P staining, and staining was also absent in sections from α-syn -/- mouse brains; attesting to the specificity of the Ser129P antibody. The background in images shown in (B-D) were cropped to highlight the brain cross-sections. **(E)** Multiple sequence alignment of α-syn from different species - the Ser87 and Ser129 sites are boxed. Note the high degree of conservation in the Ser129 site in mammals, unlike the Ser87 site, which is less conserved. **(F)** Quantification of α-syn over-expression levels in vGLUT:pHl experiments. Cultured hippocampal neurons were transduced at DIV3 by lentiviruses carrying h-α-syn constructs (WT/129A/129D) tagged at the C-terminus with mScarlett (or controls), fixed at DIV 17-21, and stained with an antibody recognizing both mouse and h-α-syn. Over-expression levels were quantified by comparing α-syn levels in boutons of neurons transduced with h-α-syn (WT/129A/129D) to those transduced with the empty vector.

**Supplementary Fig. 2:**
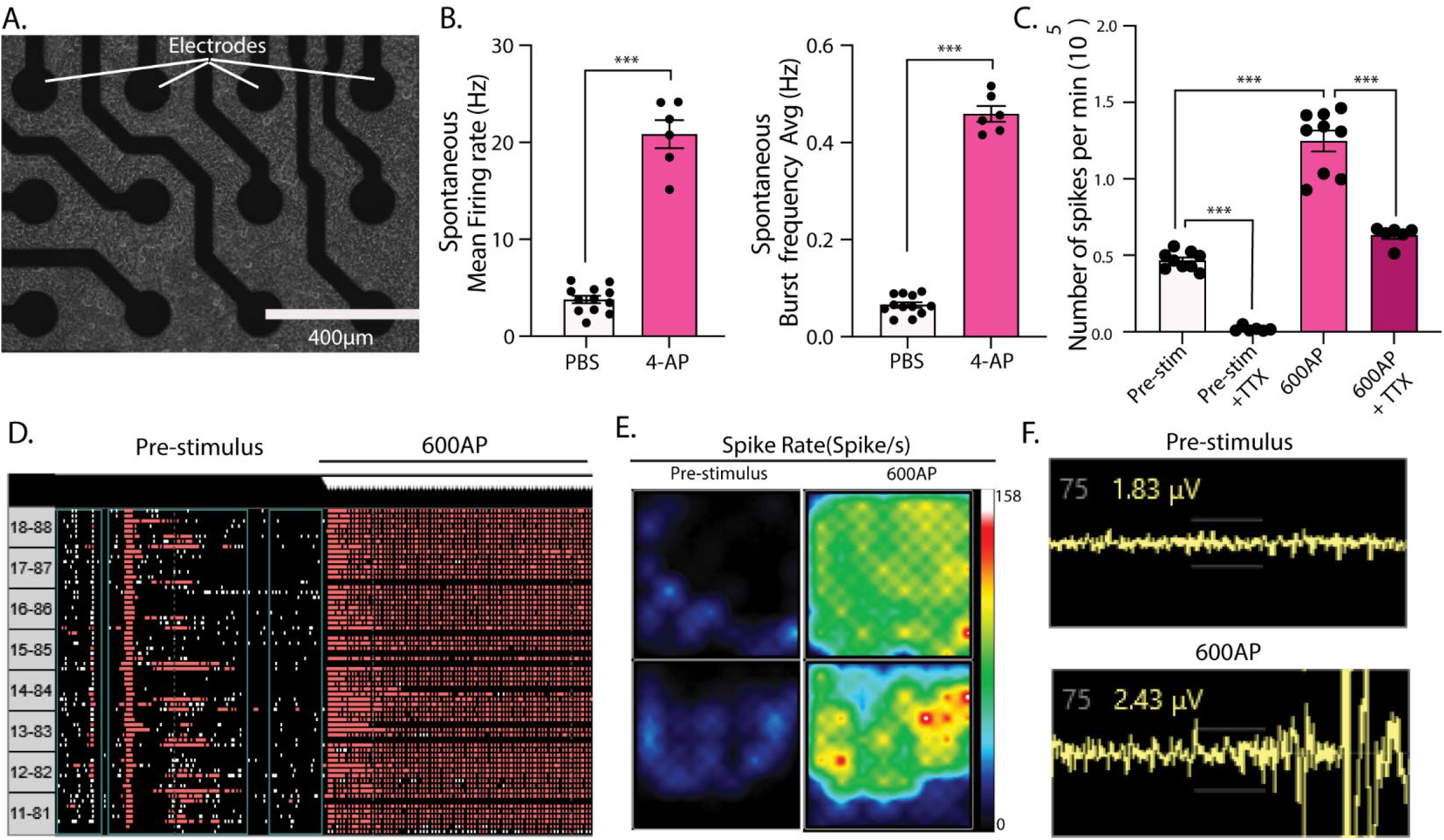
Characterization of multi-electrode array experiments. **(A)** Cultured hippocampal neurons (DIV 21) plated on a multi-electrode array. **(B)** WT hippocampal neurons were treated with 100μM 4-AP (or PBS as controls) for 3 hrs and subjected to MEA analysis of mean firing rate, burst frequency, and evoked activity using. Data shown as mean ± SEM (\*\**p* < 0.01, ***,*p* < 0.001 by Student’s *t* test; n = 6–12 independent experiments). **(C)** WT hippocampal neurons were electrically stimulated (600AP/10Hz), with or without 0.5 μM TTX (pre-incubated for 15 mins), and spike number counts were recorded. Note increased spikes after electrical stimulation, which are blocked by TTX. **(D)** Representative raster plot showing network activity pre- and post-stimulus (600AP). **(E)** Spike activity heat map visualized using Axion BioSystems Integrated Studio (AxIS). **(F)** MEA recording traces pre- and post-stimulus (600AP). Raw voltage recording of a single well from a 6-well MEA containing 64 electrodes is shown.

**Supplementary Fig. 3:**
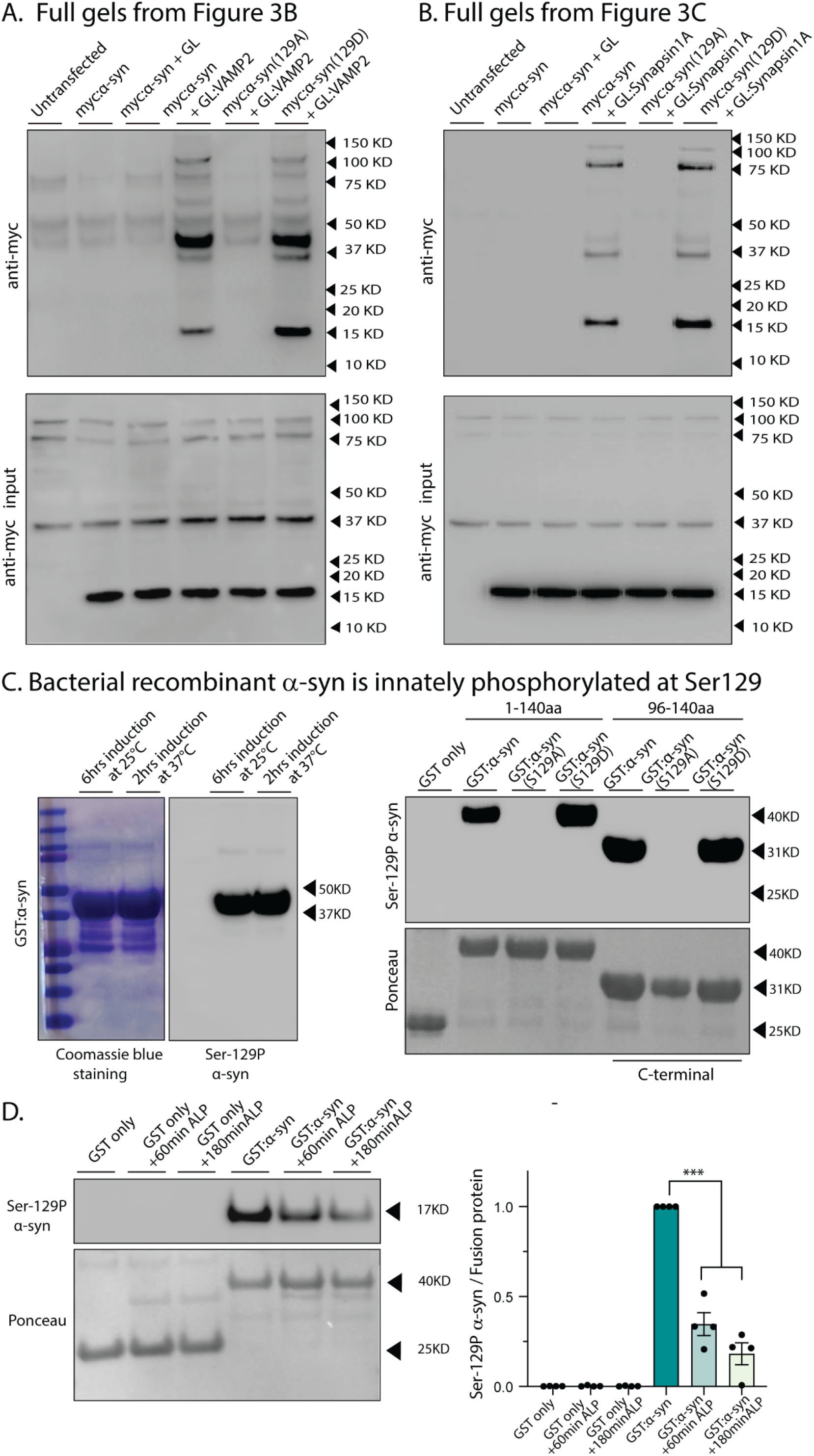
Full gels and characterization of the bacterially synthesized recombinant α-syn. **(A, B)** Full blots from co-IP experiments in neuro2a cells. Rendering the α-syn Ser129-site phospho-incompetent prevents α-syn interaction with both VAMP2 and synapsin. Note that some non-specific bands are seen with the anti-myc antibody. **(C)** GST-α-syn fusion protein expressed in bacteria is innately phosphorylated at Ser129, and two different induction protocols showed similar results. <u>Left:</u> Coomassie-blue stained NuPAGE Bis-Tris polyacrylamide gel, and immunoblotting with an α-syn Ser129P antibody. <u>Right:</u> Characterization of full-length (1-140 aa) or C-terminus (96-140 aa) α-syn GST-fusion proteins – with or without the phospho-incompetent (S129A) mutation. Samples were analyzed by NuPAGE and immunoblotted with the α-syn Ser129-P antibody (top panel). Ponceau staining (bottom panel) shows equivalent loading of fusion proteins. Note that all phospho-competent bacterially synthesized α-syn is endogenously phosphorylated at the 129-site (the phospho-incompetent 129A cannot be detected by the phospho-specific antibody). **(D)** Incubating recombinant α-syn GST fusion proteins with alkaline phosphatase (ALP) for 60-80 minutes attenuated Ser129P, confirming that the bacterially synthesized protein is endogenously phosphorylated at this site; blots quantified on right (\*\*\**p* < 0.001).

**Supplementary Fig. 4:**
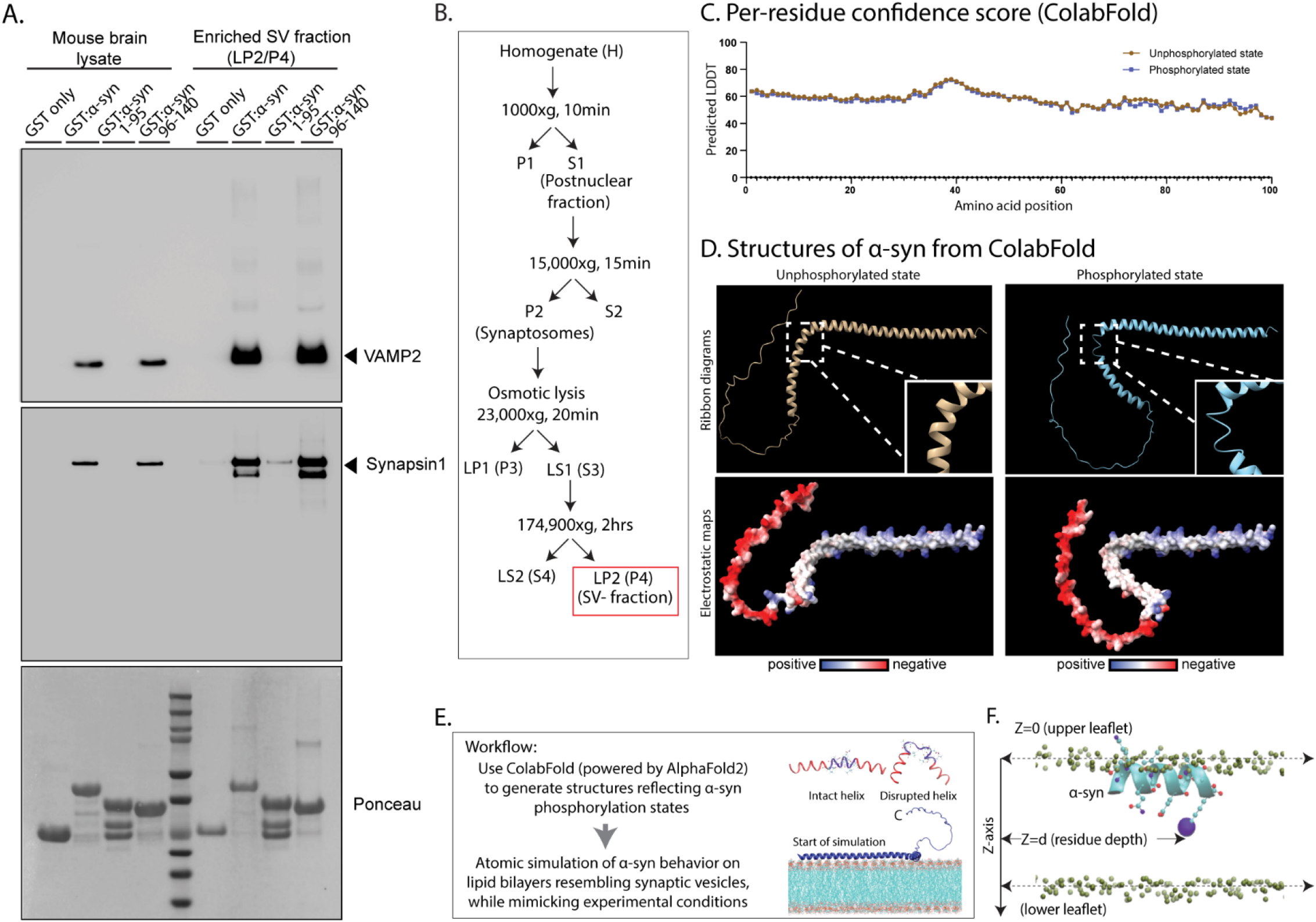
Binding-site of VAMP2 and synapsin on α-syn, fractionation assay and simulation details. **(A)** Both VAMP2 and synapsin interact with the C-terminus of α-syn. GST-tagged α-syn – full-length (1-140 aa), membrane-binding domain (1-95 aa), and C-terminal domain (96-140 aa) – was immobilized on beads and incubated with mouse whole-brain lysates or mouse SV-fractions (LP2). Material bound to the GST-α-syn (full-length and truncated) beads was immunoblotted for VAMP2 and synapsin. Note that both VAMP2 (top gel) and synapsin (middle gel) bound to full-length and C-terminus (96-140 aa) α-syn, but not to the membrane-binding domain (1-95 aa) of α-syn. Ponceau staining (bottom gel) shows equivalent loading of fusion proteins in all lanes. **(B)** Sub-fractionation protocol used in this study for isolating enriched SVs from mouse brain. The LP2 (also called P4) fraction (red box) was used for all experiments in this study. **(C)** Per-residue confidence scores – predicted local-distance difference test (plDDT, see Mariani et al., 2013) – for the ColabFold structures used in this study. **(D)** Starting ColabFold-generated structures that were used in the membrane-binding simulations. Ribbon diagrams (top) of the unphosphorylated and phosphorylated states. Insets show zoomed images of the regions where the protein loses helicity (residues from ∼ 62-70) and becomes more flexible in the phosphorylated state, exaggerating the kink at this location. Electrostatic maps (bottom) highlight the N-terminus neutral membrane-binding surface in the center (white) with positive lysine “ladder” (blue) – an arrangement thought to play a critical role in membrane binding. Note that this arrangement is largely conserved in both unphosphorylated and phosphorylated states. **(E)** Structures corresponding to unphosphorylated (permanent Ser at 129 position) and phospho-mimic (S129D) were generated using Collabfold, and membrane-binding was simulated (see text and methods). Illustrations on the right show altered helicity upon disruption of hydrogen bonds and the overall principle of the membrane-binding simulations. **(F)** Schematic showing boundary definitions of membrane leaflets and protein residue depths used to calculate membrane penetrance of α-syn; viewed from the top.

**<u>Legend for Supplementary Movie 1</u>**: Movies spanning 30-120 ns of the molecular simulations, capturing major transitions that occur during α-syn C-terminus folding – unphosphorylated state (left) and constitutively Ser-129 phosphorylated state (S129D, right). Note that initially, both scenarios start with an open structure. As the simulations progress, phosphorylated α-syn undergoes conformational changes due to crowding of five positively-charged lysine residues (60K, 80K, 96K, 97K, and 102K) around the negatively-charged phosphorylated S129 residue. In contrast, the unphosphorylated molecule largely remains in an extended confirmation over time. We speculate that the stability conferred by the phosphorylation-induced conformation changes in the C-terminus (96-140 residues) facilitate the interaction of VAMP2 and synapsin, which bind to this domain (see **Supp. Fig. 4**).

